# Generation of Functional Human T Cell Development in NOD/SCID/IL2rγ^null^ Humanized Mice Without Using Fetal Tissue: Application as a Model of HIV Infection and Persistence

**DOI:** 10.1101/2022.05.30.494009

**Authors:** Chloé Colas, Olga Volodina, Kathie Béland, Tram N.Q. Pham, Yuanyi Li, Frédéric Dallaire, Clara Soulard, William Lemieux, Aurélien B.L. Colamartino, Camille Tremblay-Laganière, Renée Dicaire, Jean Guimond, Natasha Patey, Suzanne Vobecky, Nancy Poirier, Éric A. Cohen, Elie Haddad

## Abstract

Generating humanized mice with fully functional T cells currently relies on co-implantation of hematopoietic stem cells from fetal liver and autologous fetal thymic tissue (BLT mouse). However, access to such tissues has ethical and logistical challenges. Herein, we show that NOD/SCID/IL2rγ^null^ mice humanized with cord blood-derived CD34^+^ cells and implanted in quadriceps with pediatric thymic tissues excised during cardiac surgeries (CCST mice) are an alternative to BLT mice. Our data revealed a strong immune reconstitution in CCST mice, with T cells originating from CD34^+^ progenitor cells, proliferating efficiently in response to mitogenic stimulation *ex vivo* and capable of rejecting allogeneic human leukemic cells *in vivo*. Despite having less T cells than BLT mice, CCST mice were equally susceptible to mucosal or intraperitoneal HIV infection. Importantly, HIV-specific T cell responses were significantly higher in CCST-mice (median: 10.4% *vs*. 0.7%; *p*<0.0001 for CD8^+^cells and 3.9% *vs*. 0.7%; *p*<0.01 for CD4^+^ cells). As well, antiretroviral therapy (ART) robustly suppressed viremia and reduced the frequencies of cells carrying integrated HIV DNA by up to 2 logs in various CCST mouse tissues. As in BLT mice, we observed a complete viral rebound in 67% of the animals by 2-4 weeks following ART interruption, suggesting the presence of HIV reservoirs. In conclusion, CCST mice represent an ethical and practical alternative to BLT mice, broadening the feasibility of utilizing humanized mice for research on HIV and other human diseases.

**One Sentence Summary:** We herein report a new humanized mouse model implanted with human cord blood hematopoietic stem cells and allogenic pediatric thymic tissue that develops a functional T cell compartment and supports efficient HIV infection and persistence during antiretroviral therapy.

## Introduction

For the last three decades, the development of humanized mice aiming at modelling more faithfully the human immune system has been an important quest. Indeed, several human diseases cannot be modelized in rodents, thus hindering research in immune pathophysiology and drug development. To obtain a humanized immune system, the most commonly used model is achieved by injecting human hematopoietic stem cells (HSC) in sublethally irradiated NOD/SCID/IL2rγ^null^ (NSG) mice (humNSG). This humNSG model allows for the development of partially functional lymphoid and myeloid human cells (1, 2). Indeed, in the absence of a human thymic environment, HSC-derived T cell progenitors cannot undergo full maturation and education since the mouse thymus is atrophic in NSG mice, and, in addition, does not express human leukocyte antigen (HLA) molecules (3). To address this issue the Bone-marrow-Liver-Thymus (BLT) model was developed (1, 4, 5). In this BLT model, fetal human thymic pieces are engrafted under the mouse kidney capsule, along with an i.v. injection of HSC purified from the autologous fetal liver. This model allows a more authentic thymopoiesis and consequently, displays an improved human T cell reconstitution and function compared to the humNSG (1, 4, 5). Humanized models that exhibit a multilineage immune reconstitution and a functional T cell compartment, such as the BLT, are paramount in studies of human-specific pathogens such as human immunodeficiency virus (HIV) which infects cells of the immune system.

Mice are not susceptible to HIV infection owing to the lack of essential virus-dependency host factors that HIV relies on to support viral entry, expression and production of infectious particles (6). Thus, the development of humanized mice represents a major advance in this field in that it enables studies of HIV replication, pathogenesis, immune responses and sensitivity to therapeutics (7). When inoculated with HIV, humNSG mice develop high levels of viremia with HIV invading multiple tissues (8-11). However, a caveat of the humNSG model is the absence of the adaptive human immune response and in this context, the BLT model is more superior and currently considered the most advanced and versatile small animal system to study HIV transmission, persistence and treatment (12, 13). BLT mice infected with HIV have high levels of sustained viremia, display virus-induced CD4^+^ T cell depletion and are capable of mounting virus-specific humoral and cellular immune responses (14). Importantly, viral replication in BLT mice can be completely suppressed by antiretroviral therapy (ART), making the model useful to investigate HIV persistence and to develop experimental strategies for eliminating/controlling persistent HIV reservoirs (15-18).

The BLT mouse model is extensively used to study the human immune system and is conceivably the gold standard to study HIV and human immune system interactions. However, access to human fetal tissues is challenging or even impossible in several countries, making this model not broadly available for research worldwide. Moreover, given the size of fetal tissues, the number of mice that can be made from one donor is limited. This constraint increases the variability during pre-clinical investigations, underlining the need for a larger production of mice per donor. Our goal was to develop a humanized mouse model using easy-to-access pediatric thymic tissues obtained during cardiac surgeries (thymectomy is often performed during major cardiac surgeries) that could support an optimal T cell reconstitution when co-engrafted with human cord-blood derived HSC. Herein, we report that this Cord blood Cardiac Surgery Thymus (CCST) humanized mouse model exhibits a functional T cell compartment that derives from CD34^+^ cells and recapitulates key features of HIV infection and persistence.

## Results

### Histological Structure of Primary and Secondary Lymphoid Organs Recovered from BLT and CCST Mice

The CCST model is based on the engraftment of pediatric thymic tissues, otherwise discarded in the context of cardiac surgery, into quadriceps femoris muscle of the mouse (Fig. 1A-B), similar to what is being done in human thymus transplantation (19, 20). We first assessed the need to culture thymic pieces *in vitro* (as shown in Figure 1-A) prior to engraftment in order to eliminate mature T cells. All mice (n=6) engrafted with uncultured thymic pieces died within two months from what appeared to be a graft-versus-host-disease (GvHD)-like syndrome (Supp. Fig 1). In contrast, most mice (5/6) engrafted with previously cultured thymic pieces, as performed in human thymus transplantation (19, 20), survived at least 20 weeks post-humanization (p=0.0006, Supp. Fig. 1).

**Figure 1.**
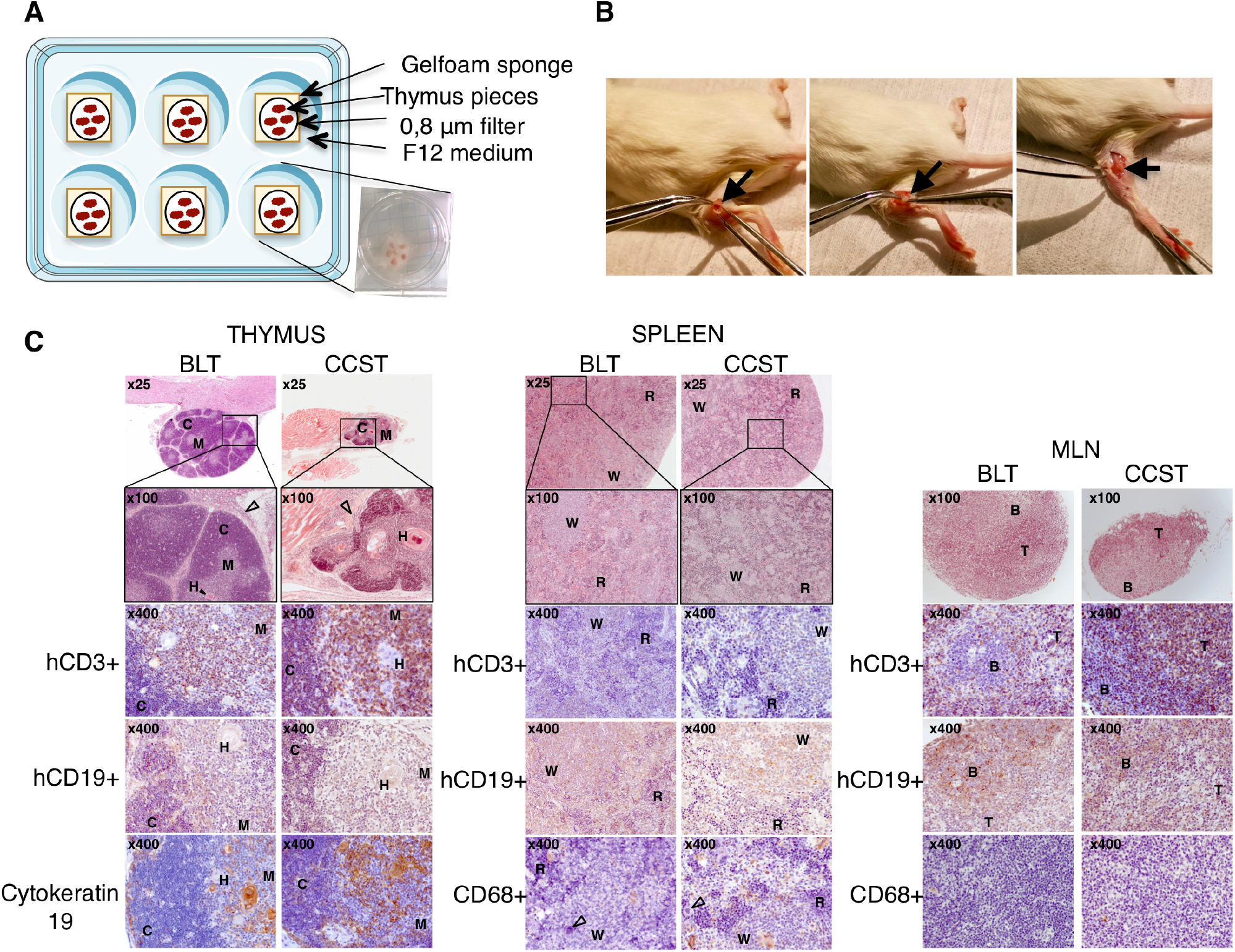
Pediatric thymus from cardiac surgery can be implanted into NSG mice as an alternative to fetal thymus. **A)** Thymus retrieved from 3-day to 6-year-old donors undergoing cardiac surgery was cut to 1mm^2^ pieces and put on a gelfoam sponge culture system for 7 to 21 days. **B)** 2-3 pieces were engrafted into the quadriceps muscle of NSG mice along with an i.v. injection of 1-2×10^5^ umbilical cord blood CD34^+^ (see supporting video in supplemental). **C)** Representative thymic tissue sections from both BLT and CCST mice recovered at 30 weeks post-humanization (Left panels). Hematoxylin-phloxin-safran staining shows that engrafted tissues were well inserted in the surrounding tissues (arrowhead, in the kidney for BLT; in the quadriceps muscle in CCST). Both models showed typical thymus structure with cortex -C-medulla -M-and Hassall’s corpuscles -H-. Immunochemistry staining shows the presence of hCD3^+^ (T cells) and hCD19^+^ (B cells). Thymus displayed medullar and cortical arrangement with thymic epithelium expressing cytokeratin 19. Spleen tissues from both BLT and CCST (Middle panels) displaying white - W-and red -R-pulp, T cells (hCD3^+^), B cells (hCD19^+^) and monocytes (hCD68^+^), as well as megakaryocyte (empty arrowhead). Mesenteric lymph nodes (MLN) recovered from BLT and CCST mice (Right panels) displayed germinal centers with B and T lymphocyte zones.

Fetal thymic pieces engrafted under the kidney capsule of BLT mice were capable of growing to more than 10 times their size. In contrast, CS thymus from CCST mice did not exhibit a size expansion and was indeed difficult to recover and separate from the muscle fiber. Nevertheless, histological analysis showed that in both cases the thymi were well integrated in the surrounding tissues with the presence of a connective tissue (Fig. 1C). Moreover, in both models, thymi were similarly populated with numerous CD3^+^ cells and some CD19^+^ cells. Hassall’s-like corpuscle structures were observed (Fig. 1C-Left panel) in both models, although larger in the CSST mice, indicating an increased activity of the thymus. Unlike in mice humanized without a human thymus graft (humNSG), mesenteric lymph nodes (MLN) with organized structure, including germinal center with distinct T and B lymphocyte zones, were developed in both BLT and CCST mice (Fig. 1C-Right panel). As well, spleens with comparable structures were observed in both models (Fig. 1C-Middle panel). Hence, CCST mice developed primary and secondary lymphoid organs that were similar to those observed in the BLT model.

### Human Immune Cell Reconstitution in CCST Mice

The kinetics of human immune cell reconstitution was assessed in peripheral blood. In BLT and CCST models, human CD45^+^ cells could be observed as early as two weeks after humanization (Fig. 2A). All three models showed comparable kinetic and intensity of hCD45^+^ reconstitution (*p*=0.4228, two-way ANOVA). Indeed, BLT showed a reconstitution going from 56% ± 19% to 73% ± 26% of hCD45^+^ cells between 10-and 20-weeks post-humanization (wph) while CCST and humNSG both plateaued at approximately 60% by 5 wph (Fig. 2A-upper graphs, 64% ± 24 at 10 wph and 56% ± 33% at 20 wph for CSST; 46% ± 5.2 at 10 wph and 64% ± 2% at 20 wph for humNSG). Significant differences between groups were observed for hCD3^+^ (calculated among hCD45^+^-Fig. 2A) (*p*<0.0001 for BLT vs CCST; *p*<0.0001 for BLT vs humNSG; *p*<0.001 for CCST vs humNSG; Two-way ANOVA). While only less than 1% of hCD3^+^ cells were detected at 10 weeks post-humanization in humNSG, BLT mice supported the development of human T cells within 2 to 3 weeks post humanization, resulting in a rapid and robust reconstitution of hCD3^+^ population (16% ± 24%; Fig. 2A). As in BLT mice, CCST mice displayed early T cell reconstitution, with the detection of hCD3^+^ cells after 2 weeks (3.6% ± 4.8%), which then increased gradually throughout the life of the mice (Fig. 2A). Regarding hCD19^+^, although the shape of the reconstitution curve was similar in all three models, significant differences in the proportion of hCD19^+^ among hCD45^+^ were observed between the groups (*p*<0.0001 for BLT *vs* CCST; *p*<0.0001 for BLT *vs* humNSG; *p*<0.01 for CCST *vs* humNSG, two-way ANOVA). No significant differences were found for hCD14 population (*p*=0.8420, two-way ANOVA).

**Figure 2.**
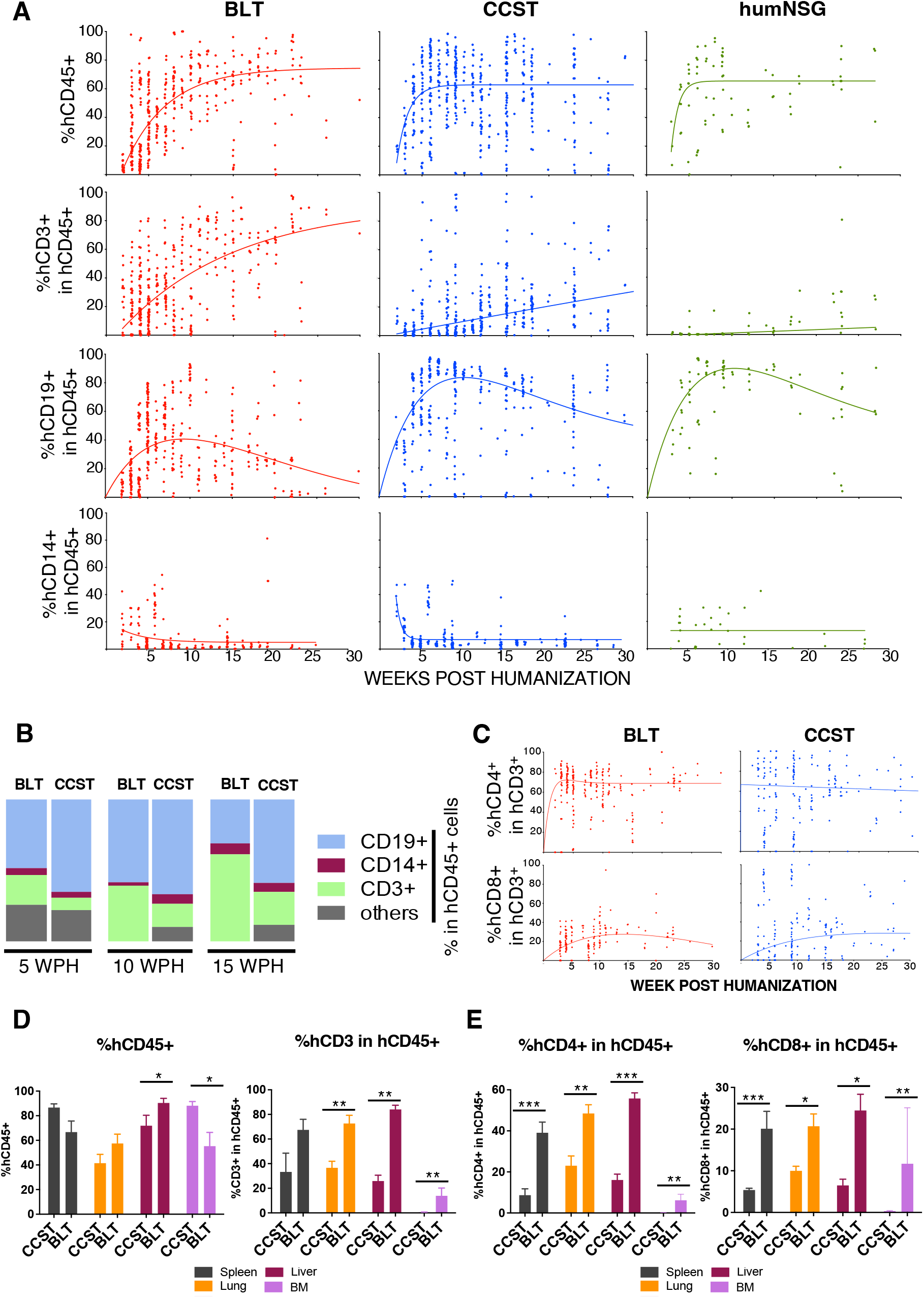
Unlike humanized mice without a thymus (humNSG), CCST and BLT humanized models allow for robust T cell reconstitution. **A**. Peripheral blood from mice of the three models was harvested and analyzed by flow cytometry for the level of human cell reconstitution (each dot represents a mouse analyzed at a given time point). Extent of the reconstitution of hCD45^+^ (calculated among total CD45^+^ cells (mouse and human)), hCD3^+^ (among hCD45^+^), hCD19^+^ (among hCD45^+^) and hCD14^+^ (among hCD45^+^) was as shown. Non-linear regression curve was calculated for each subpopulation, using the growth exponential association model for all populations except hCD19, for which a two-phase exponential association model was applied. **B**. Graphic representation of the relative proportions of human T (CD3^+^), B (CD19^+^) and monocytic (CD14^+^) cells in BLT and CCST mice at 5, 10, and 15 weeks post-humanization showing an increase in T cell proportion through time for BLT mice. **C**. Dynamic changes in the level of human CD4^+^ and CD8^+^ T-cells within the hCD3^+^ population in peripheral blood of CCST and BLT mice. Non-linear regression curves were calculated, using a two-phase exponential association model. **D**. Proportions of human CD45^+^, CD3^+^ and (**E**) levels of CD4^+^ and CD8^+^ T-cells in tissues of CCST and BLT mice at 25 weeks post-humanization. Mann-Whitney’s test was applied to compare ranks of two groups; *p<0.05, **p<0.01, ***p<0.001, ****p<0.0001.

The relative proportion of the different human immune cell populations in the blood was compared between BLT and CCST mice. In BLT mice, the hCD3^+^ population increased in proportion through time and stabilized with human cells composed of mainly T cells (hCD3^+^), followed by B cells (hCD19^+^) and monocytes (hCD14^+^) (Fig. 2B). In contrast, CCST had predominantly hCD19^+^, balanced with hCD3^+^ and hCD14^+^ as well as other hCD45^+^ cells (Fig. 2B). Interestingly, this cell distribution remained stable through time. Within the hCD3^+^ population, the proportions of hCD4^+^ and hCD8^+^ cell populations were comparable between BLT and CCST mice (*p*=0.6500 for hCD4 between models, and *p*=0.0995 for hCD8, two-way ANOVA), although CCST data showed a higher variability between mice (Fig. 2C). Analysis of the human cell immune population in the lung, liver, bone marrow and spleen showed that levels of hCD45^+^ cells were overall comparable between the two models, while that of hCD3^+^ were significantly higher in tissues of BLT mice compared to CCST (Fig. 2D). Consequently, human CD4^+^ and hCD8^+^ proportions among the human immune cells (hCD45^+^) were also significantly higher in tissues of BLT mice relative to CCST mice (Fig. 2E). Altogether, the CCST model reconstituted a human immune system that was qualitatively and quantitatively an intermediate between BLT and humNSG mice, with a robust hCD3^+^ compartment and secondary lymphoid organs, two key characteristics for immune function.

One could argue that T cells observed in CCST mice are thymus-derived proliferating mature T cells, mimicking the model where NSG mice are injected with PBMC. However, this is highly unlikely for several reasons. First, the kinetics of T cell count rise in CCST was slow and as such, could not have been a result of proliferation of peripheral mature T cells (Fig 2A). Second, in the absence of cord blood CD34^+^ cells, NSG mice engrafted with a cardiac surgery thymus, which had been cultured under the same condition as in CCST mice, had little circulating T cells (< 1%, not shown). Nevertheless, to formally rule out this possibility and confirm that T cells in CCST mice actually stemmed from CD34^+^ cells that seeded the thymus and differentiated into thymic progenitors and mature T cells, CCST mice were engrafted with CD34^+^ cells previously transduced with a GFP-expressing lentivirus. First, from the apparition of hCD3 until week 7 post-humanization, our data suggest that there was a transient mix of T cells originating from CD34^+^ HSC and of T cells coming directly from the CS thymus. Then, after week 7, based on GFP^+^ hCD3^+^ proportion, these T cells originating from the implanted thymus seemed to have been replaced by the engrafted CD34^+^ cells. Indeed, after 7 weeks post-humanization, we observed that the percentage of GFP^+^ cells was similar in all lineages (B, T or CD14^+^), strongly suggesting that, from this point on, T cells did originate from the CD34^+^ cell pools (Fig 3A). Interestingly, the same was true for BLT mice as illustrated in Fig 3B.

**Figure 3.**
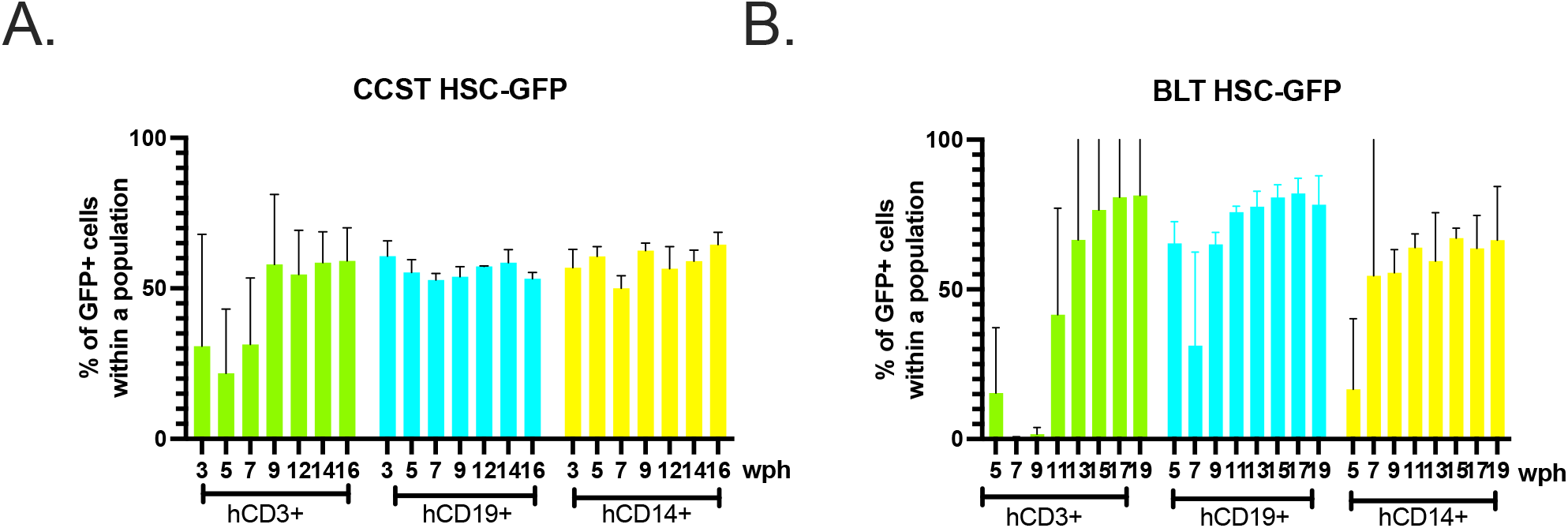
Circulating human immune cells in CCST and BLT mice are originating from engrafted CD34^+^hematopoetic stem cells. Human HSCs were transduced with a GFP-expressing lentivirus. Transduction efficiency was 39% at the time of mouse engraftment. Shown are immune reconstitution data from (A) CCST (n=3) and (B) BLT (n=2) mice up to 19 wph. T cell (hCD3^+^), B cell (hCD19^+^) and monocyte (CD14^+^) compartments exhibited similar proportions of GFP-expressing cells. Y axis is indicative of the percentage of GFP+ cells within the population indicated in the x axis (either hCD3+, hCD19+ or hCD14+).

### T cell Function in CCST and BLT Mice

We tested T cell function using both an allogenic human tumor challenge and an *ex vivo* phytohemagglutinin (PHA)-dependent proliferation assay. The tumor challenge was performed by injecting allogeneic luciferase-expressing human pre-B leukemic cell lines REH (Fig. 4A-C). All REH-challenged humNSG reached high levels of luciferase expression 3 to 6 weeks following tumor cell injection and died within 7 weeks, while no REH leukemic cells were detected in any of the BLT mice, as shown by the absence of luciferase signal and 100% survival rate (Fig. 4A-C). In comparison, the response to REH challenge was more variable with CCST mice. Indeed, 5 out of the 14 CCST mice (35%) injected with REH controlled leukemic cell growth while the other CCST mice allowed for the establishment of leukemia, but with a significant delay in the kinetic of leukemic cell dissemination, and consequently in the survival (*p*<0.001, Fig. 4 A-C), compared to humNSG (p=0.0002). Despite this relative control of leukemic growth, as a whole, CCST mice had a higher luciferase activity (p=0.0111) and poorer survival than BLT mice (p=0.0135).

**Figure 4.**
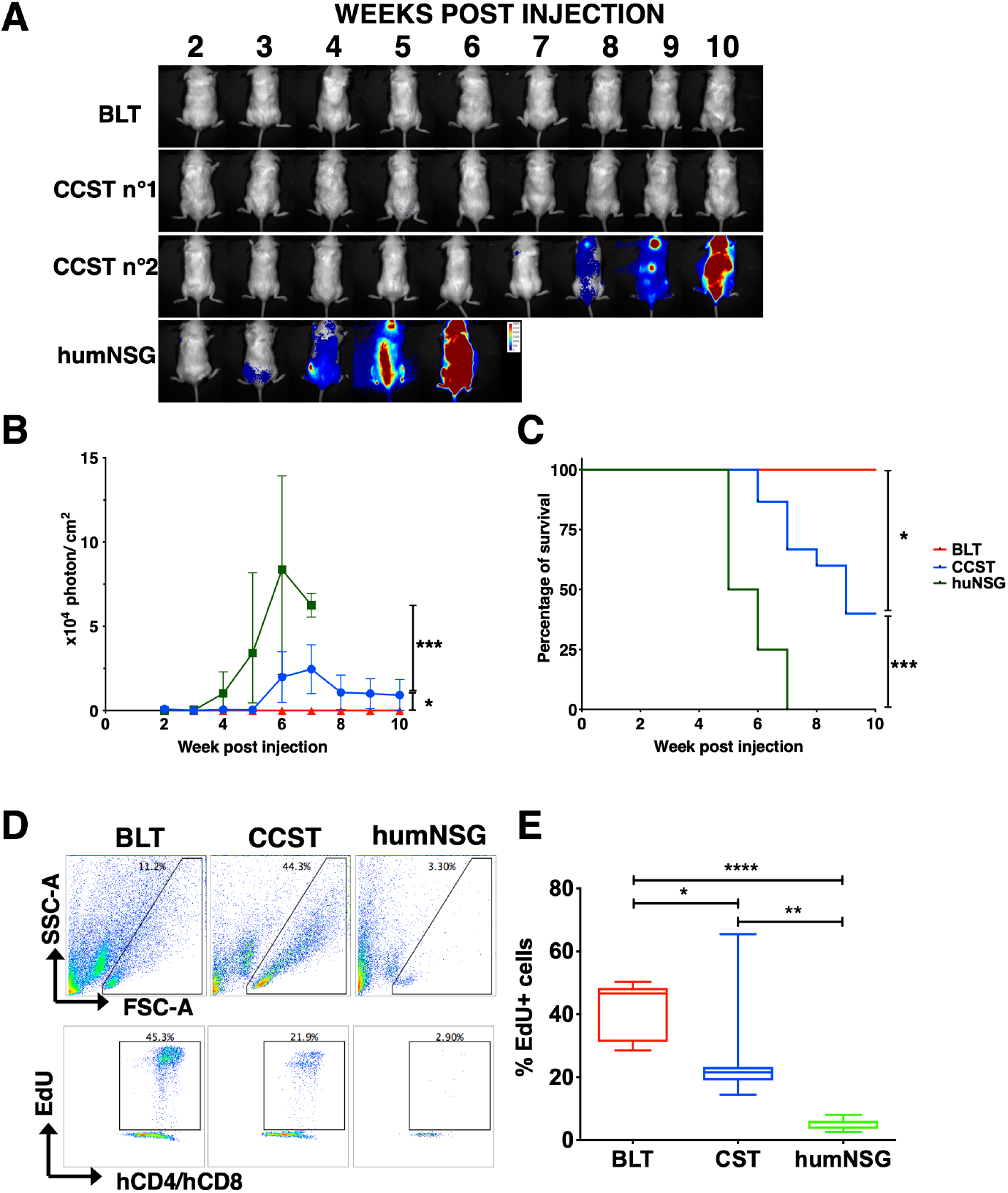
Functional T cells are developed in CCST and BLT mice. **A**. Pre-B leukemic REH cells expressing luciferase were injected in the three humanized models and mice were imaged weekly by injecting D-luciferin to follow the progression of the leukemic cells. Depicted are representative BLT, CCST and humNSG mice with different capability to respond to the REH challenge. **B**. Intensity of the luciferase expression (expressed in photon per cm^2^) was used as a surrogate marker of leukemia progression in BLT mice (red line), CCST mice (blue line) and humNSG mice (green line). Medians +/-range are shown for the luciferase signal, p=0.0111 between BLT and CCST, p=0.0002 between humNSG and CCST, and p<0.0001 between BLT and humNSG: two-way ANOVA. **C**. Survival curves show a significant difference (Log-rank tests) between BLT (n=10, red line), CCST (n=14, blue line) and humNSG (n = 8, green line). BLT vs humNSG: p<0.0001; BLT vs CCST: p=0.0135; CCST vs humNSG: p=0.0001. **D**. Proliferative capacity of peripheral blood T cells isolated from the three models of humanized mice was tested by *ex vivo* stimulation with 5 μg/ml of PHA-L for 24 hours in the presence of EdU. Representative flow cytometry plots are shown for each model. **E**. Enumeration of hCD4^+^hCD8^+^Edu^+^ cells showed that T cells from CCST (n=7) and BLT (n=8) mice proliferated significantly more than humNSG (n=7) mice (p=0.0061 with CCST, p<0.0001 with BLT). T cells isolated from CCST mice proliferated also less than BLT (p=0.0466). One-way ANOVA with Tukey post-test. *p<0.05, **p<0.01, ***p<0.001, ****p<0.0001, n.s. = not significant.

In the context of PHA-dependent proliferation assay, both hCD4^+^ and hCD8^+^ T cells from CCST and BLT mice incorporated EDU (5-ethynyl-2’-deoxyuridine) following mitogen stimulation, albeit at a lower level for CCST mice (median of 46.67% for BLT and of 21.62% for CCST, *p*=0.0466 one-way ANOVA with Tukey post-hoc test). In contrast, T cells isolated from humNSG did not respond to PHA stimulation (Fig. 4D&E, median of 5.69%, *p*=0.0061 compared to CCST, *p*<0.0001 compared to BLT, One-way ANOVA with Tukey post-hoc test). T cells isolated from BLT and CCST mice had respectively a mean of 8 and 5-fold more EdU^+^ cells than those from humNSG (Fig. 4E, *p*<0.0001, one-way ANOVA). Altogether, these data indicate that T cells from CCST mice were functionally capable of proliferating upon mitogen stimulation and rejecting tumor cell challenge.

### Robustness of the CCST model

In order to assess whether it was possible to generate CCST mice with biobanked tissues, we compared animals made with fresh thymic tissues to those made with previously frozen thymic pieces. As shown in Supplemental Figure 2, mice humanized with biobanked tissues were comparable to those generated with fresh tissues in terms of the level of CD3^+^ cells and T cell ability to reject infused leukemic cells. Overall, the success rate was 74.8% (242 mice; 27 groups-defined by mice made on the same day with the same tissues and HSC). Mice within groups were widely homogenous in that they were either all usable or not at all, except for 3 of the 27 groups in which there was a mix in the quality of the immune reconstitution, suggesting that the variability principally lied between the groups and not within a group. Finally, the impact of HLA compatibility between thymic pieces and injected CD34^+^ cells on the quality of immune reconstitution was assessed in 103 CCST mice and no significant impact was detected (*p*=0.9232, Chi-Square test, Supp. Table 1). Similarly, the age of the engrafted thymus did not affect significantly the level of immune reconstitution (*p*=0.0732, Chi-Square test, Supp. Fig. 3A) nor T cell function, as evaluated by the ability to clear REH (p=0.4805, two-sided Chi-Square test, Supp. Fig. 3B&C).

### CCST Mice Support Efficient HIV Infection and Mount Robust HIV-Specific T Cell Responses

As stated above, BLT mice are the gold standard of humanized mouse models to study pathogenesis and treatment of HIV infection (12, 13). We thus compared key characteristics of HIV infection in CCST and BLT models using HIV NL4.3-ADA-GFP (Fig. 5A). First, we observed that both models were susceptible to HIV infection via mucosal or intraperitoneal route, as viral loads were detectable at similar levels through time post-infection (Fig. 5B). As well, real-time PCR analysis showed nearly comparable abundance of total viral DNA in different tissues although frequencies of integrated HIV DNA-positive cells in spleen, lung and liver were significantly higher in BLT mice (Fig. 5C). Interestingly, in CCST mice, frequencies of productively infected (p24^+^) CD4^+^ T cells was higher in the liver although they were not significantly different in spleen or lung tissues. (Fig. 5D; gating strategy illustrated in Supp. Fig. 4).

**Figure 5.**
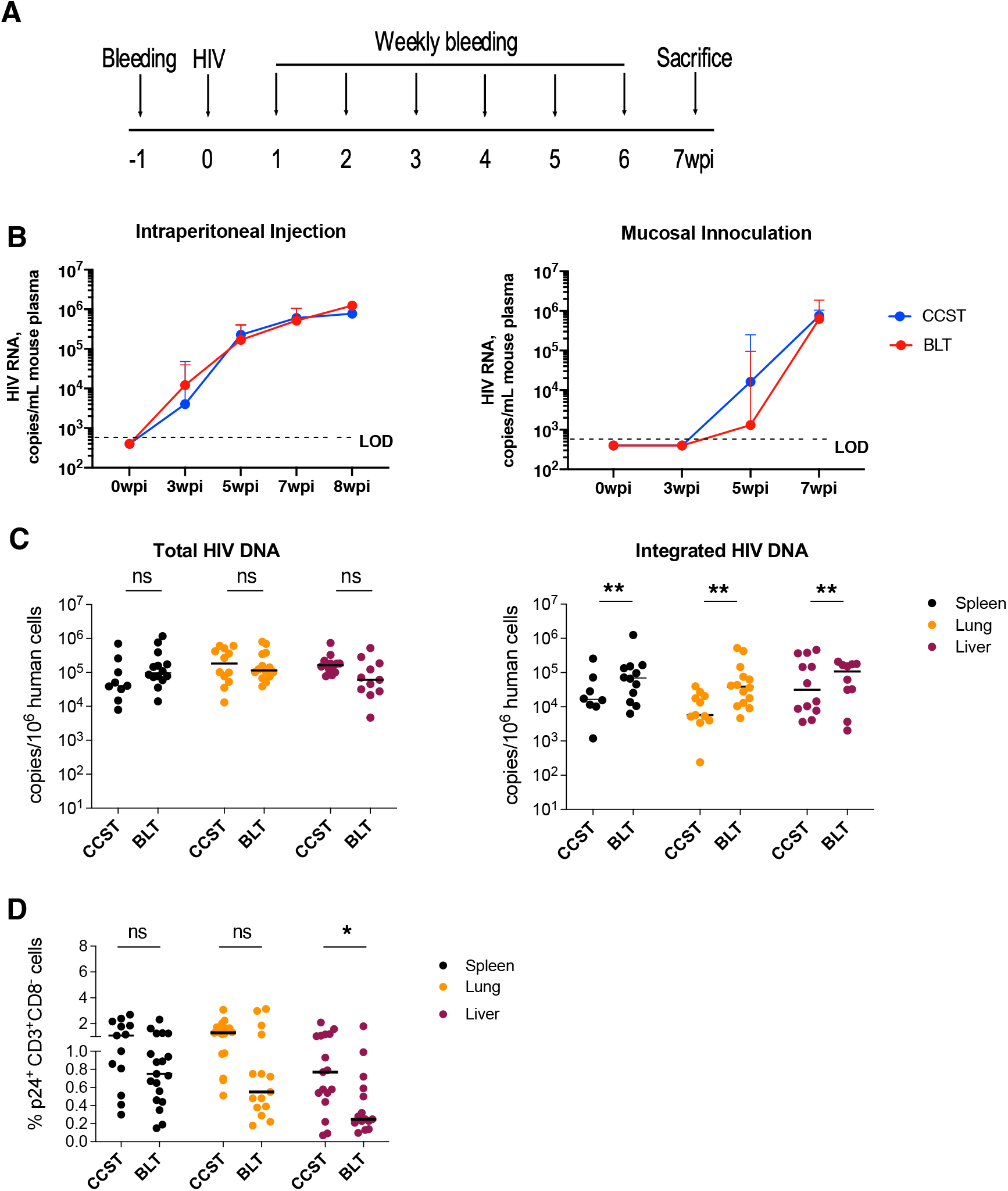
CCST mice are as susceptible to HIV infection as BLT mice A. CCST and BLT mice were infected with 200,000 TCID_50_ of NL4.3-ADA-GFP HIV via intraperitoneal injection or vaginal inoculation and monitored for 7 weeks. **B**. Plasma viral load measured in CCST (blue lines) and BLT (red lines) mice following HIV infection. Shown are median values (+95% confidence interval) from 9 to 19 (depending on the time point) CCST and 15 BLT mice injected intraperitoneally and 7 CCST and 6 BLT mice inoculated vaginally. **C**. Levels of total and integrated HIV DNA in the spleen, lung and liver, quantified by real-time PCR. **D**. Frequency of virus-expressing (p24^+^) human CD4^+^ T-cells (as defined by hCD3^+^hCD8^-^ owing to HIV-mediated CD4 downregulation) in the spleen, lung and liver as measured by flow cytometry. Mann-Whitney’s test was applied to compare ranks of two groups. ns, not significant; wpi, weeks post infection. LOD - limit of detection.

Having demonstrated that CCST mice were highly susceptible to HIV infection, we next asked whether T cells from infected animals could elicit T cell response upon *ex vivo* stimulation with HIV peptides. For this, we pulsed T cells from the spleen with HIV peptide pools (21) or PMA and then assessed their capacity to produce IFNγ. To this end, we observed that in response to HIV peptides, significantly higher levels of CD4^+^ and CD8^+^ T cells in CCST mice were producing IFNγ compared to BLT mice (median: 10.4% *vs*. 0.7%; *p*<0.0001 for CD8^+^ T cells and 3.9% *vs*. 0.7%; *p*<0.01 for CD4^+^ T cells, respectively p < 0.0001; Fig 6A and Suppl Fig 5). As well, PMA stimulation resulted in more CD4^+^ T cells from CCST mice expressing IFNγ although no differences were noted for CD8^+^ T cells. A similar analysis of splenic T cells from uninfected CCST and BLT mice showed that in response to mitogenic stimulation with PMA/ionomycin, both CD4^+^ and CD8^+^ T cell subsets were able to produce IFNγ, highlighting once again that T cells developed in CCST mice are functional. However, we found no statistically significant difference in the frequency of IFNγ-expressing T cells between CCST and BLT mice (Fig. 6B) although T cells from the spleen of CCST mice expressed a higher level of activation markers HLA-DR and CD69 relative to those from BLT mice (Fig. 6C).

**Figure 6.**
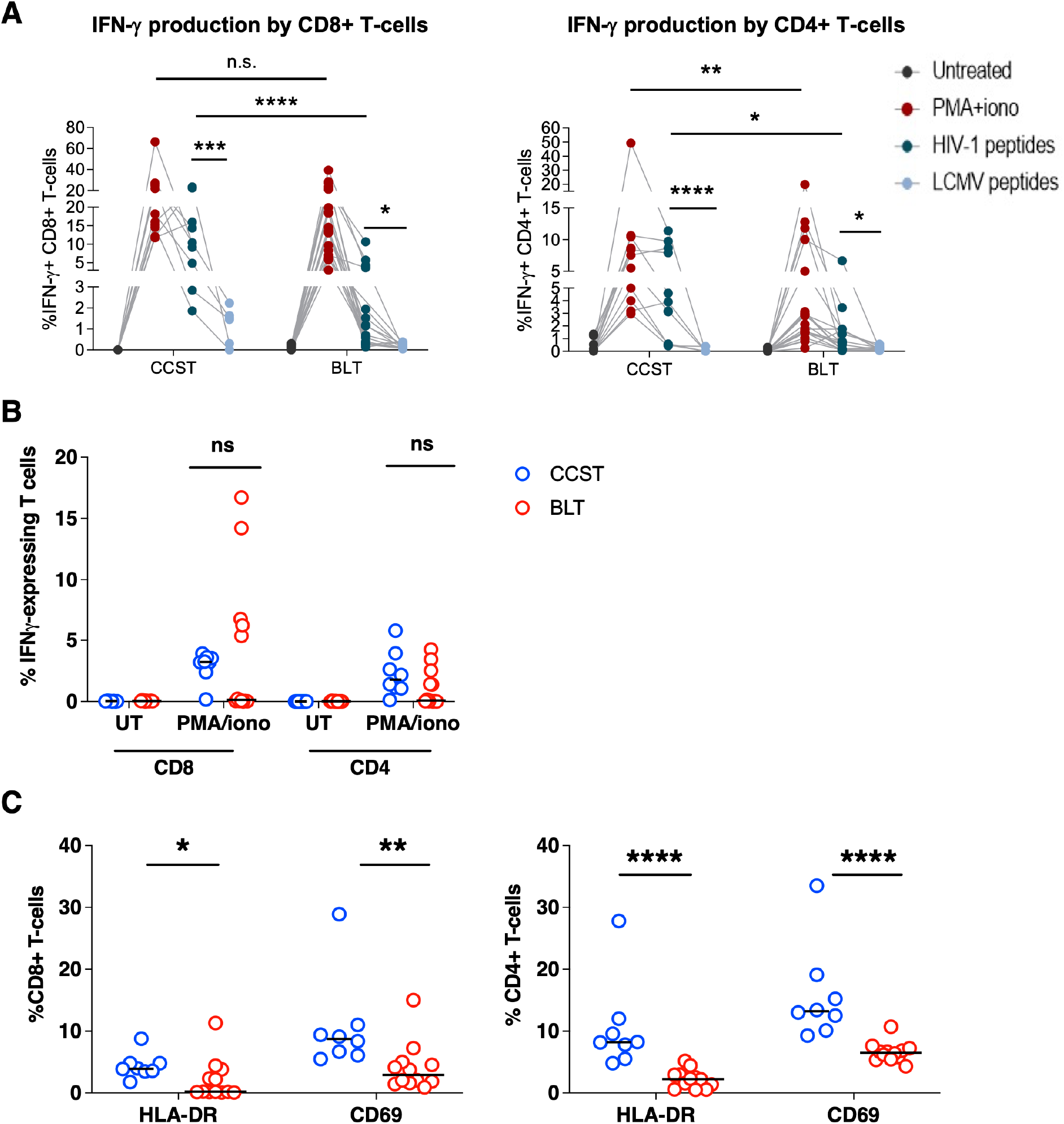
Comparison of general and HIV-specific T cell response in CCST and BLT mice. **A**. Splenocytes from infected CCST and BLT mice were kept untreated or stimulated with PMA/ionomycin, pooled Clade B HIV peptides (env, gag, pol, nef) (2 μg/mL) or LCMV peptides (GP61-80, GP276-286) (100 ng/mL). LCMV stimulation served as a specificity control for the assay. Frequencies of cytokine-expressing human Tcells were measured intracellularly by flow cytometry. Depicted are overall data in different mice (CCST, 10 mice and BLT, 12 mice). Four dots connecting each line represent an individual mouse. **B**. Splenocytes from uninfected (UI) CCST and BLT mice were stimulated with PMA/ionomycin and assessed for IFN-γ expression in T cells as described in Panel A. **C**. Surface expression of activation markers HLA-DR and CD69 on T cells from the spleen of uninfected CCST (blue) and BLT (red) mice were measured by flow cytometry. In all panels, each dot is one mouse. Mann-Whitney’s test was applied to compare ranks of two groups; *p<0.05, **p<0.01, ***p<0.001, ****p<0.0001, n.s. = not significant.

Overall, our findings revealed that CCST mice could be efficiently infected with HIV and that T cells developed in this model were functional, capable of expressing cytokines following antigenic stimulation.

### CCST Mice as an Investigative Model to Study HIV Persistence during ART

Having shown that CCST mice were capable of supporting efficient infection and mounting an antigen-specific T cell response, we next wanted to establish whether this humanized mouse model could be used to characterize the development of HIV reservoirs. To study this aspect, we infected CCST and BLT mice and when the level of viremia started plateauing (about 6-7 weeks post-infection), the animals were treated with an antiviral regimen containing emtricitabine, tenofovir and raltegravir (Fig. 7A). As expected, plasma viral loads were reduced to undetectable levels in ART-treated mice in both models (Fig. 7B). Congruent with this observation, we observed a significant reduction in the level of total (5-24 folds) and integrated (12-33 folds) HIV DNA in different lymphoid and non-lymphoid tissues (Fig. 7C) in ART-treated mice. As well, ART resulted in a marked median reduction of cell-associated HIV RNA by at least 45 to 77 folds depending on organs (Fig. 7D). Lastly, the proportion of p24^+^hCD3^+^hCD8^-^ cells in tissues was suppressed to nearly undetectable levels in ART-treated mice in both models (Fig. 7E, *p*<0.001 in all ART conditions). Importantly, the fact that we observed a complete viral rebound upon treatment interruption demonstrates not only the presence of HIV reservoirs in infected CCST mice (Fig 7F) but also that CCST mice can be used as a model of HIV persistence during ART.

**Figure 7.**
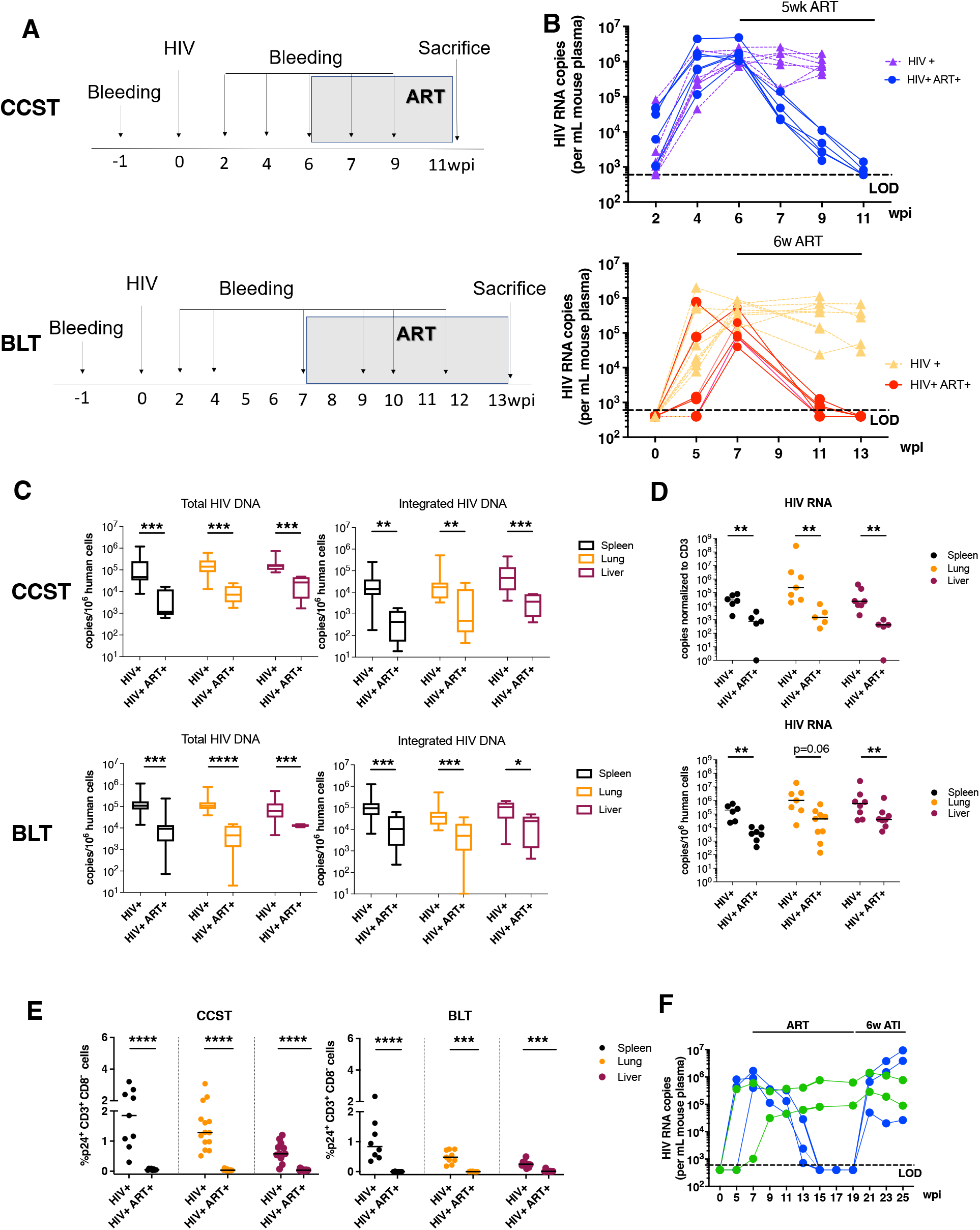
Establishment of HIV latency in the CCST mouse model. **A**. CCST mice were infected (i.p route) with 200,000 TCID_50_ of NL4.3-ADA-GFP HIV. At 5-6 wpi, a group of infected mice was treated for up to six weeks with emtricitabine (100 mg/kg weight), tenofovir (50 mg/kg) and raltegravir (68 mg/kg). Control group was represented by untreated mice. **B**. Dynamic changes in plasma viral load over the course of infection and ART. Untreated mice are depicted in purple (CCST) and yellow (BLT), ART-treated mice in blue (CCST) and red (BLT). **C**. ART-induced changes in levels of total and integrated HIV DNA in different tissues of CCST and BLT mice. **D**. HIV RNA levels in tissues of untreated or ART-treated CCST and BLT mice. **E**. Effect of ART on the frequency of CD4^+^ T cells in tissues expressing viral protein p24. **F**. Presence of viral rebound following antiretroviral therapy treatment interruption (ATI-blue lines), supporting the existence of HIV latency in infected CCST mice. As a result of ATI, levels of viremia were increased to the level of untreated mice (green lines). Each line represents one mouse. Mann-Whitney’s test was applied to compare ranks of two groups; *p<0.05, **p<0.01, ***p<0.001, ****p<0.0001, n.s. = not significant.

## Discussion

Herein, we describe a new model of humanized mice that fosters the development of functional T cells without the use of fetal tissues. This model comes in a timely fashion since research projects conducted with fetal tissues from abortions are becoming increasingly criticized. Indeed, abortion of pregnancies beyond 12 weeks of gestation is prohibited in many countries, and in 2019 the US Department of Health and Human Services announced that the National Institutes of Health would be highly restrictive on research using fetal tissues(22). Therefore, there is a need for the development of new humanized mouse models that could substitute the BLT model without losing its advantages. in this study, we provide strong evidence that CCST mice could be a good alternative to BLT mice, especially for HIV research. Indeed, CCST displayed strong immune reconstitution, with T cells originating from CD34^+^ progenitor cells, proliferating efficiently in response to mitogenic stimulation *ex vivo* and capable of rejecting allogeneic human leukemic cells *in vivo*. These findings demonstrate the functionality of developing human T cells in CCST mice. Despite having less T cells than BLT mice, CCST mice were equally susceptible to mucosal or intraperitoneal HIV infection and mounted a stronger HIV-specific T cell responses following infection. Importantly, the presence of integrated HIV DNA could be documented despite ART treatment and virological suppression, and viral rebound could be observed upon treatment interruption, indicating that CCST mice can be used as a model to study HIV persistence in the presence of antiretroviral therapy.

The CCST model is different from the recently developed, NeoThy humanized mice (23) in two major ways. In the latter model, neonatal thymic pieces of approximately 10 days old neonates were implanted under the kidney capsule along with i.v. injection of CD34^+^ cells isolated from cord blood. Ours involves implantation of thymic tissues, which could be up to 5 years old, in the quadriceps instead of kidney capsula. We used this approach since initial attempts at implanting thymic pieces in the kidney capsule were repeatedly unsuccessful (data not shown), prompting us to ultimately adopt the procedure used for thymic transplantation in human (19, 20) in which the thymus is implanted in the quadriceps. According to our veterinary surgeon, the surgical procedure is less complex. Another advantage of this technique is that we could freeze thymic pieces and cord blood CD34^+^ cells to perform humanization when needed (Fig. Supp. 2). Interestingly, as described in thymic transplantation in human (24), and as observed for the NeoThy model (23), HLA-compatibility did not have an impact on the reconstitution of the immune system (Supp. Table. 1). However, the use of pediatric thymi seems to require a pre-culture *ex vivo* to prevent rapid occurrence of a GvHD-like syndrome. In this context, we hypothesize that resident T lymphocytes from the thymic graft are capable of expanding and attacking recipient tissues and cells, leading to mouse death. As such, the pre-culture step is essential to eliminate potentially xenoreactive mature/maturing T cells from the engrafted thymus. Consequently, T cells developed in CCST mice, when the humanization is stabilized (after 7 weeks), are derived from the injected HSC that are niched in the humanized mouse bone marrow, as demonstrated by the results from our GFP-transduced HSC experiments. The thymic culture could explain the differences observed in the reconstitution kinetic of hCD3^+^ cells in CCST mice compared to the BLT, in which a pre-culture of fetal tissues is not needed. Kalscheuer et al published a personalized model of immune-mediated disorders based on the engraftment of fetal thymus along with stem cells from adult patients. Interestingly, in this model, it is necessary to administer anti-CD2 antibodies to eliminate mature T cells that might otherwise cause GvHD (5). This suggests that *ex vivo* culture or monoclonal antibody administration to eliminate mature T cells from thymic pieces reduces the occurrence of GvHD. Moreover, the T cell immune reconstitution observed in our CCST mice is similar to the one described in the NeoThy model, even though we injected a lower number of CD34^+^ cells, suggesting that the CCST model is not dependent on the amount of engrafted CD34^+^ as observed in the NeoThy model (5).

The CCST humanized mice displayed organized primary and secondary lymphoid organs, suggesting the development of a functional immune system. The implanted thymus in CCST mice exhibited medullar and cortical arrangement with classical thymic epithelium expressing cytokeratin 19 as in the case with BLT mice. As well, the spleen from CCST mice shared similar structures with megakaryocytes observed in both models, indicating splenic extramedullary hematopoiesis. Lastly, mesenteric lymph node displayed germinal centers having distinct T and B lymphocyte zones in both the BLT and CCST models.

The immune reconstitution in the peripheral blood of CCST mice was robust, albeit not as high as in the BLT, but clearly more efficient than humNSG, particularly with respect to the T cell compartment. Interestingly, the replenishment of T cells in the CCST model was progressive, as observed after T cell-depleted bone marrow transplantation in human (25). Similar to the BLT system, a transient first wave of T cells likely comprises mature T cells coming directly from the engrafted thymus, however, as humanisation stabilises, peripheral T cells observed in CCST mice are T cells stemming from HSC that undergo a physiological differentiation in the transplanted thymus. This observation is supported by the similar distribution of GFP-expressing cells within hCD3, hCD19 and hCD14 cells. It is very unlikely that CD34^+^ cells differentiate in mouse thymus in the CCST given that we observe a good human thymic maturation with Hassal corpuscules and that humNSG mice do not display the same kinetic of T cell development. Moreover, the relative proportions of immune subpopulations remained stable through time in CCST mice. As previously reported (4), the reconstitution of human CD45^+^ cells was slightly slower in BLT mice, but ultimately reached higher levels. This difference in the reconstitution kinetics might be partially due to the source of hCD34^+^ (fetal liver *vs*. cord blood), different number of injected cells (5 × 10^5^ *vs*. 1-2 × 10^5^, respectively) and xenogeneic T cell proliferation.

Importantly, the CCST model recapitulated key T cell functions such as PHA-induced proliferation and allogenic tumor rejection. This approach could be useful to study immune interactions with tumors, as one could model patient-derived xenograft (PDX) and implant autologous immune system from the same patient’s HSC together with an allogenic thymus from cardiac surgery (either fresh or biobanked).

The classical humNSG model does not require the use of fetal tissues. Nevertheless, in the context of HIV research, the lack of a functional adaptive immune system in this model limits its capacity to help address important questions pertinent to virus-host interactions. Humanized NSG mice are also less susceptible to HIV infection through the mucosal route, which represents the main mode of viral transmission in human.(26) In this study, we demonstrate that CCST mice represent an ethical and efficient substitute for the BLT model for HIV study. First, we show that CCST mice are equally susceptible to HIV infection through both the mucosal and intraperitoneal routes. Despite having fewer CD4^+^ T cells as targets for HIV infection, productive viral dissemination in CCST animals was very efficient across different organs relative to BLT mice, with higher frequency of infected cells actively expressing viral proteins. Second, we provide evidence showing that viral replication in CCST mice could be efficiently suppressed by ART and that there is *bona fide* viral rebound upon treatment interruption. The latter strongly supports the existence of authentic HIV reservoirs in these infected mice. Third, we demonstrate that HIV exposure leads to the development of a more vigorous HIV-specific T cell response in CCST mice compared to the BLT. In this setting, CCST mice represent the first humanized animals generated without the need of fetal tissues and yet, capable of eliciting antigen-driven immune responses, especially in the context of HIV infection.

Altogether, this new model of mouse humanization is robust and convenient by: (1) its wide range of potential thymus donors, (2) tissue abundance allowing for larger production, (3) the use of biobanked tissues and (4) HLA independence. However, the CCST model has its own limitations. First, there is a greater variability compared to the BLT in terms of the overall reconstitution, relative distribution of immune subpopulations and success rate (74.8%). The different distribution of immune cell subsets (Fig 2) might be linked in part to the technical difficulty in determining whether the thymic pieces to be implanted were components of the medulla and/or cortex. Since fetal thymus is smaller than pediatric thymus, pieces of the latter are more likely to comprise both medulla and cortex than the former ones, which could explain the greater variability observed in the CCST model. However, although the level of T cells in remains lower compared to BLT, this limitation does not constraint the levels of HIV viremia nor the extent of viral infection.

In conclusion, the CCST model is an ethical and practical alternative to the BLT model that could prove to be better for HIV studies because of its robust antigen-specific T cell response. The ease by which CCST model can be achieved, in regard to surgery skills, availability of thymic material, as well as the fact that thymic sections can be cryopreserved and biobanked, could dramatically increase the availability of functional humanized mouse models worldwide and facilitate, among others, *in vivo* HIV research.

## Materials and Methods

### Study approval

The study was reviewed and approved by the Centre Hospitalier Universitaire (CHU) Sainte-Justine institutional (CER#2126) and the CSSS Jeanne-Mance review boards (Montreal, Canada), and was performed in accordance with federal and provincial laws. Human tissues were obtained following written informed consent to participate in this study. All animal experiments were performed in accordance with protocols approved by each institution’s Institutional Animal Care and Use Committee (IRCM 2015-11 & 2018-06 and CIBPAR#582&593) following Good Laboratory Practices for Animal Research.

### Tissue processing

Human autologous fetal thymus and liver were harvested and processed on the same day. Fetal liver was cut into small pieces and shaken at 230 rpm for 30 min at 37°C in a sterile complete RPMI medium (RPMI 1640+GlutaMAX™ (Gibco™ by Life Technologies™, Thermo Fisher Scientific, Waltham, MA) + 10% decomplemented fetal bovine serum (FBS) (Gibco™ by Life Technologies™) and Penicillin-Streptomycin (P/S) (Wisent, St-Jean Baptiste, QC, Canada) supplemented with 1mg/ml collagenase (Sigma Aldrich, St-Louis, MO), DNAse I recombinant (Roche by Sigma Aldrich) and sodium pyruvate (Sigma Aldrich). The suspension was then filtered with a 70 μm cell strainer and washed with Dulbecco’s phosphate buffered saline without calcium or magnesium (dPBS^-/-^) (Gibco™ by Life Technologies™) with 0.6% citrate phosphate dextrose (Sigma Aldrich). Hepatocytes were then removed by low-speed centrifugation (18G, 5 min) and the supernatant spun again (470 x *g*, 5 min). Pelleted cells were collected and purified by density centrifugation using FICOLL (GE Healthcare, Uppsala, Sweden). HSC were then enriched by using hCD34 MicroBead kit Ultrapure (Miltenyi, Bergisch Gladbach, Germany). hCD34^+^ cells were either resuspended in dPBS^-/-^ and kept on ice until injection in mouse or were frozen in FBS + 10% dimethyl sulfoxide (DMSO) (Thermo Fisher Scientific).

Human pediatric thymus tissues were obtained from patients undergoing cardiac surgery (n=19, age ranges between 1 day to 5 years old; mean age: 1 year and 2 months, median: 6 months) during their care at CHU Sainte-Justine (Montreal, Canada). Upon reception, thymus was washed in dPBS^-/-^ and cut into small pieces (around 4-8mm^3^). Pieces were either put in culture or frozen in decomplemented human serum type AB (Wisent, Canada) +10% DMSO (10-20 pieces/ml) as described (5). When put in culture, pieces were placed on top of a 0.8-um isopore membrane filter (Sigma Aldrich), on top of a 1-cm^2^ absorbable gelatin sponge (Pfizer, Kirkland, QC, Canada) in the presence of F12 medium (Gibco™ by Life Technologies™) + 10% FBS + P/S + Fungizone (Gibco™ by Life Technologies™) + HEPES (Sigma Aldrich) (Fig. 1A). Thymic pieces were kept in culture at 37°C between 7 to 21 days. Fetal thymus was either used directly for surgery in mice or frozen as described above.

### Cord blood CD34^+^ isolation

Human cord blood was obtained from the Cord Blood Research Bank of the CHU Sainte-Justine under the approval of the CHU Sainte-Justine institutional review board and written informed consent from donors. Mononuclear cells were purified by using SepMate™ (Stemcell technologies, Cambridge, MA, USA) using FICOLL. As described above, HSC were then enriched by using hCD34 MicroBead kit Ultrapure. hCD34^+^ cells were either kept in culture at 37°C in RPMI medium with human recombinant Stem Cell Factor (SCF) at 50 ng/ml (PeproTech, Rocky Hill, NJ, USA), human recombinant thrombopoietin (TPO) (PeproTech), human recombinant Fms-related tyrosine kinase 3 ligand (FLT3-L) (PeproTech) until injection in mice (within 3 to 6 hours of culture) or were frozen in FBS + 10% DMSO.

### Human tissue transplantation

NOD/LtSz-scid IL-2Rγc(null) (NSG) mice were purchased from Jackson Laboratory (Bar Harbor, ME, USA). All mice were housed in specific pathogen-free animal facility of CHU Sainte-Justine Research Center or in the animal core facility of the Montreal Clinical Research Institute. For all models, six-to ten-week-old NSG mice were conditioned with sublethal (2.5 Gy) total-body irradiation a day prior to humanisation. HumNSG mice were simply injected i.v. with 1×10^5^ of pooled human cord blood CD34^+^ cells in 100 μl of dPBS. For BLT mice, human fetal thymus fragments (1 per mouse) measuring about 1mm^3^ were implanted underneath the recipient kidney capsule. Within 24 hours, 3-5×10^5^ human autologous fetal liver CD34^+^ cells in 100 μl of dPBS^-/-^ were injected intravenously. For CCST mice, three human pediactric thymus fragments were implanted in the left quadriceps muscle of the mouse and 1-2×10^5^ pooled human cord blood CD34^+^ cells in 100 μl of dPBS were injected intravenously. When mentioned, CD34^+^ were transduced with a lentivirus coding for GFP. Briefly, 1×10^5^ CD34^+^ cells isolated from cord blood were plated on RetroNectin-coated wells and transduced with lentiviral particles based on baboon envelope pseudotyped (BaEV-LV) (27) coding for the GFP under the control of the UCOE0.7-SFFV promoter (pHUS-GFP vector) at a MOI of 1.4.(28) After 24 hours, cells were harvested, pooled, washed and injected to mice. All animals were anesthetized with isofluorane and treated with Buprenorphine for post-operative pain management and their drinking water was supplemented with Baytril antibiotic for 10 days post-surgery.

### Engraftment assessment

The presence of circulating human cells was then evaluated weekly in 50 μl of peripheral blood harvested through the saphenous vein and collected in heparinized tubes. The absolute number and proportion of cells subpopulation was assessed by FACS using CountBright™ Absolute Counting Beads (ThermoFisher) along with monoclonal antibodies directed against hCD45 (clone HI30), hCD3 (clone UCHT1), hCD19 (clone HIB19), and hCD14 (clone B159) (all from BioLegend, San Diego, CA, USA) and mCD45 (clone 30-F11) (BD Biosciences, Franklin Lake, NJ). Human CD45 percentages are calculated among all CD45^+^ cells (murine and human). Acquisition was done on a BD FACSCanto or FACS LSR Fortessa system (BD Biosciences). Mice were sacrificed when pre-defined limit points were reached. At the time of sacrifice, thymus (if present), liver, mesenchymal lymph node (if present), blood and spleen were harvested and evaluated by histopathology and flow cytometry.

### Allogeneic leukemic cell line challenge

Acute lymphocytic leukemia pre-B cells REH (ATTC®, Manassas, VA, USA) were modified to express luciferase using a pHUS Luciferase-GFP vector (constructed from the pHUS-GFP plasmid backbone and encoding for the firefly luciferase, an internal ribosome entry site (IRES) and eGFP). These cells were injected intravenously to mice (5,000 cells in 100 μl of dPBS^-/-^). Mice were then monitored weekly by injecting 3 mg D-luciferin (Perkin Elmer, Waltham, MA, USA) intraperitoneally and imaging using an *in vivo* bioluminescence imaging system (LabeoTech, Montreal, QC, Canada) system. Images were analyzed using ImageJ (version 1.52p, NIH) to quantify the bioluminescence intensity.

### T cell proliferation assay

T cell capacity to proliferate upon stimulation with phytohemagglutinin-L (PHA-L) (Sigma Aldrich) was measured following guidelines of the Click-iT® EdU cell proliferation assays kit (Thermo Fisher Scientific). Briefly, 150-200 μl of peripheral blood was drawn from the saphenous vein and collected in heparinized tubes. Red blood cells were then lysed in a buffer containing 0.15 M ammonium chloride, 10 mM potassium bicarbonate and 0.1 mM EDTA (prepared in distilled water) and white blood cells cultured with PHA-L (5 μg/mL) for 48 hours and then EdU (10 μM) for overnight. The following day, cells were harvested, stained with FITC hCD8a (clone RPA-T8, BioLegend) and hCD4 (clone RPA-T4, BD Biosciences), and then fixed/permeabilized using Click-iTTM fixative and saponin-based reagent and stained using the fluorescent dye picolylazide to reveal the incorporated EdU. Fixed cells were analyzed on the BD FACS Canto or FACS LSRFortessa system.

### Histology

The specimens were fixed in buffered-formalin phosphate 10 % and embedded in paraffin. 4 μm sections were prepared, mounted on microscope slides, and stained with hematoxylin phloxin safran. Immunohistochemistry staining was also performed on paraffin-embedded slices on the Autostainer Link 48 from Dako with the “ready to use antibodies” hCD3 (polyclonal rabbit), hCD68 (clone KP1), human Cytokeratin 19 (clone AE1/AE3) and hCD19 (EPR5906; Abcam, Cambridge, UK).

### HIV Virus production

5×10^6^ HEK 293T cells were transfected with 30 µg of CCR5-tropic pNL4.3.ADA.GFP proviral DNA (29) by the calcium phosphate method. Culture supernatant was collected at 60 h post-transfection. Virus was concentrated by ultracentrifugation over a 20% sucrose cushion and tittered in HeLa TZMbl and CEM-CCR5 cell lines (NIH AIDS reagent program). TCID_50_ was calculated using the Spearman-Karber method.

### Infection of mice and quantification of viral loads

CCST and BLT mice were infected with 200,000 TCID_50_ of NL4.3-ADA-GFP HIV in 100 µl of DMEM via intraperitoneal injection or vaginal inoculation and monitored for up to 25 weeks depending on experiments. HIV viral load in plasma of humanized mice was determined weekly using the quantitative COBAS AmpliPrep/COBAS TaqMan HIV test, version 2.0, Roche (detection limit, <20 copies/ml).

*ART*. Six-week long daily ART, consisting of emtricitabine (100 mg/kg weight), tenofovir (50 mg/kg) and raltegravir (68 mg/kg), was initiated at usually 6-7 wpi for a group of mice and maintained until the sacrifice. Control group was represented by untreated mice.

### Nucleic acid extraction and quantification of HIV genomes by real-time PCR

RNA from spleen cells or other tissues were extracted using RNeasy Mini plus kit (QIAGEN) according to manufacturer’s instructions. cDNA was generated using Superscript II reverse transcriptase (Invitrogen) and used as templates for HIV RNA analysis as detailed below. DNA from spleen cells, lung and liver cells were extracted from gDNA eliminator columns using Qiamp Fast DNA stool mini kit (QIAGEN) according to manufacturer’s instructions. Quantification of total HIV DNA, integrated HIV DNA and unspliced HIV RNA was performed by modified nested real-time PCR assay using Taq DNA polymerase (BioLabs) in the first PCR and TaqMax Fast Advanced Master Mix (Applied Biosystems) in the second PCR (30). DNA from serially diluted ACH2 cells (NIH AIDS Reagent Program, NIAID, NIH), which contain a single copy of the integrated HIV genome, was extracted and amplified in parallel to generate a standard curve from which unknown samples were enumerated. Human CD3 gene was used as a normalizer.

### Ex vivo IFN-γ production

Splenocytes from CCST and BLT mice were stimulated with PMA (50 ng/mL) plus ionomycin (500 ng/mL), or LCMV peptides (GP61-80, GP276-286) (100 ng/mL) for 1 hour or with pooled Clade B HIV peptides (env, gag, pol, nef) (2 μg/mL) (NIH AIDS Reagent Program) for overnight. Subsequently, Brefeldin-A (BioLegend) was added and cells cultured for another 6 hours. Untreated splenocytes were used as control. Frequencies of IFN*γ*-expressing human T cells were measured by flow cytometry as described below. Splenocytes from uninfected mice and LCMV stimulation were used to ensure the specificity of the assay.

### Flow cytometry

Blood cells and single-cell suspensions from tissues were stained with a combination of fluorescently labeled Abs. Dead cells were excluded using live/dead fixable violet dead cell stain kit (ThermoFisher). Surface-stained cells were fixed and permeabilized using the Cytofix/Cytoperm kit (BD Biosciences) as per manufacturer’s instructions and intracellularly stained with anti-p24 antibody or anti-IFN*γ*. Flow cytometry data were collected on a Fortessa flow cytometer (BD Bioscience) and analyzed by Flowjo software (Versions 9.9.3 and 10.1).

### Statistics

Data analysis was performed using GraphPad PRISM 8.0 (GraphPad Software). Statistical tests applied for each experiment are stated in the legends of Figures. A P value of less than 0.05 was considered statistically significant. The following symbols *, **, ***, **** signify <0.05, <0.01, <0.001, <0.0001, respectively. No statistical methods were used to predetermine population size. Randomization was not used.

## Supporting information

Suuplemental Video 1

## Supplementary Materials

Movie S1. Movie showing surgery in mouse quadriceps.

## Acknowledgments

Authors would like to thank Romas Geleziunas at Gilead Sciences for the gift of emtricitabine and tenofovir and Daria Hazuda at Merck for providing us with the raltegravir. We are also grateful to the CHU Sainte-Justine Cord Blood for Research purpose Biobank. We also thank Jaspreet Jain, Mélanie Laporte and Oussama Meziane for their experimental support. We also thank the animal and flow cytometry core facilities of the IRCM for their expertise and experimental support.

## Funding

This work was supported through funding from the Fondation Charles Bruneau, a “Chaire de Recherche Banque de Montreal” to E.H. as well as by a grant supporting the - Canadian HIV Cure Enterprise (CanCURE) from the Canadian Institutes of Health Research (CIHR) (HB2 – 164064 to E.A.C. W.L. is supported by a “Fonds de Recherche en Santé du Québec” (FRQS) scholarship award and A.C. by a Cole Foundation scholarship award. C.T-L is supported by a CIHR scholarship award. EAC is recipient of the IRCM-Université de Montréal Chair of Excellence in HIV research.

## Author contributions

C.C., Y.L., C.S., W.L., A.C., C.T-L performed the experiments pertaining to immune reconstitution and function. O.V., T.N.Q.P., and F.D. performed experiments and data analysis related to the HIV work in hu-mice. R.D. recruited participants and collected samples. J.G, N. Patey, S.V. and N. Poirier provided and processed the human tissues. C.C, O.V., K.B and T.N.Q.P. wrote the manuscript. EAC and EH conceptualized the study and wrote the manuscript. E.H. generated the hypotheses.

## Competing interests

The authors declare no potential conflicts of interest.

## Data and materials availability

Not applicable

## SUPPLEMENTAL FIGURES

**Supplemental Figure 1.**
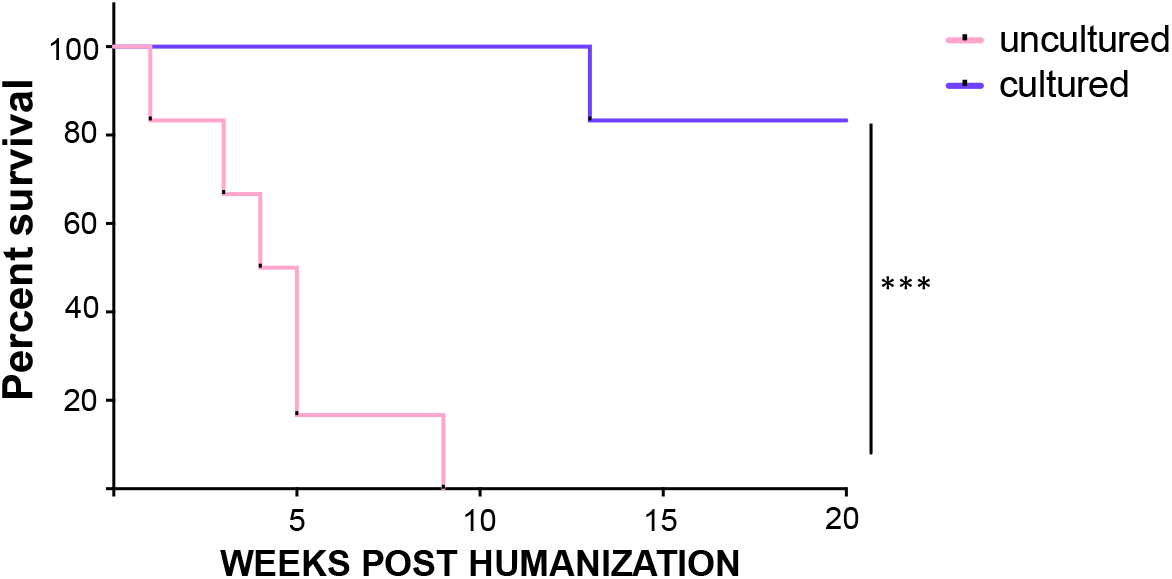
Ex vivo Culture of Thymus Prior to Engraftment Improves Survival of Recipient Mice. Human pediatric thymus obtained from cardiac surgery was cut into small pieces and either implanted directly in the quadriceps muscle of sublethally irradiated-NSG mice humanized with CD34^+^ from cord blood (uncultured, *n*=6) or put in culture (as described in Figure 1) for 7 days before implantation in the quadriceps muscle of humanized NSG mice (cultured, *n*=6). All mice that received uncultured thymus died within 9 weeks from severe GvHD symptoms, while all mice except one that received cultured thymus survived. These conditions were used in all subsequent experiments. Log-rank (Mantel-Cox) test, ***p=0.0006.

**Supplemental Figure 2.**
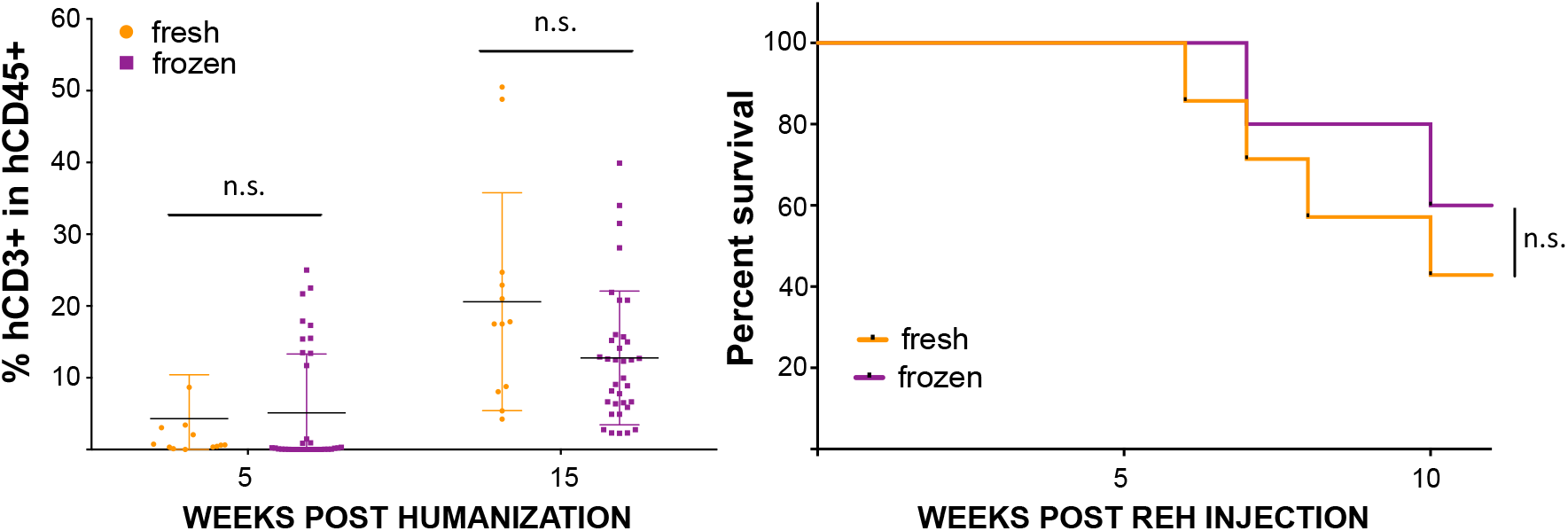
Comparison of the different treatments used before transplanting a cardiac surgery thymus into a mouse. **A**. Following dissection, cardiac surgery thymus was put in culture for immediate use or cryopreserved for future use. CCST mice made with fresh or frozen tissues were not significantly different in the level of hCD3^+^hCD45^+^ cells (left panel, T-tests) or their ability to control REH leukemia challenge (right panel, Log-rank (Mantel-Cox) test). Each dot corresponds to a mouse. N.s: not significant.

**Supplemental Figure 3.**
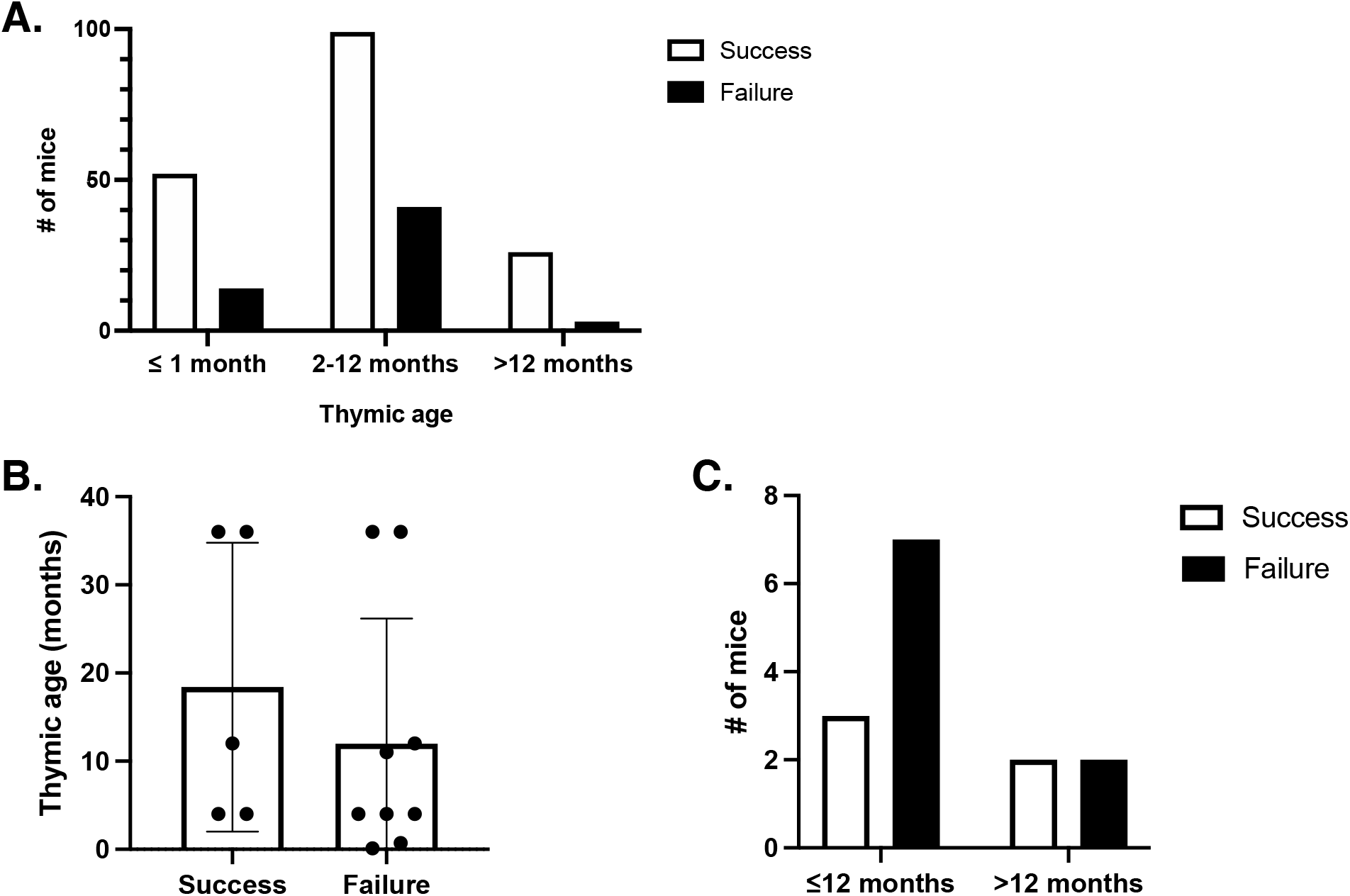
Impact of thymic age on the success of immune reconstitution and clearance of tumor cells. **A. Success (**n=177) or failure (n=58) of mice in reconstituting an immune system with early T cell output according to the age of the engrafted thymus. The data show that thymic age did not have an effect on the immune reconstitution (p=0.0732, Chi-Square test). Success was defined by early T cell output in immune reconstitution and functional assays (when available). **B**. Mean (+/-SD) of thymic age according to REH clearance (Success group, n=5) or death of the animal (failure group, n=9), p=0.4567, unpaired T-test. **C**. Effect of thymic age on mouse surviving REH challenge. p=0.4805, Two-sided Chi-Square test.

**Supplemental Figure 4.**
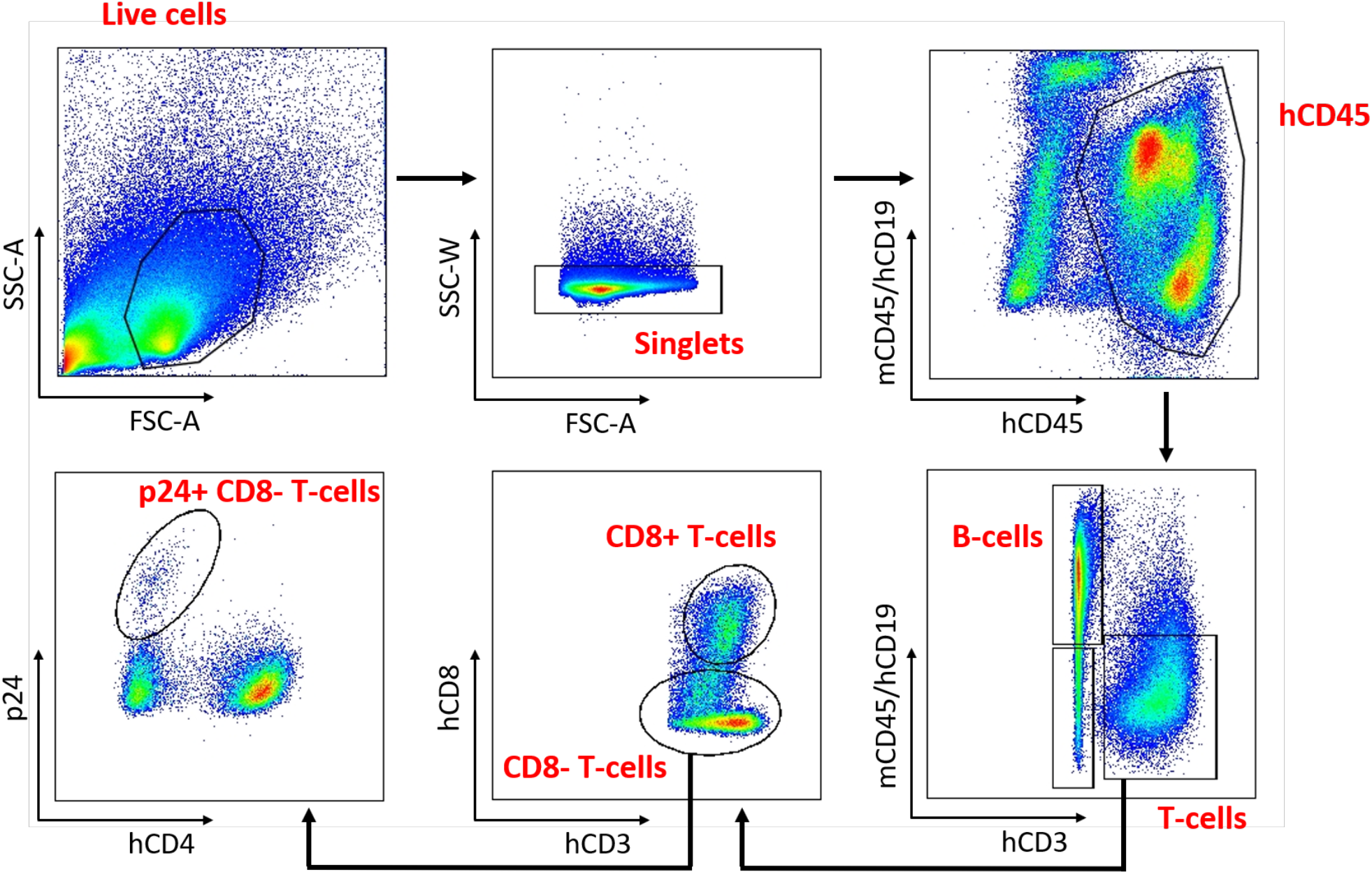
Gating strategy for flow cytometry analysis to identify human immune cell subsets, including HIV-infected T-cells. Single cells were isolated from peripheral blood or tissues and stained with a mixture of fluorescent-labeled antibodies according to the cell surface staining protocol. Total human immune cell population was identified using anti-human CD45 antibodies and further characterized as B-cells (hCD45^+^CD19^+^), monocytes (hCD45^+^CD14^+^), T-cells (hCD45^+^CD3^+^) and their subtypes, CD8^+^ T-cells (hCD45^+^CD3^+^CD8^+^) and CD4^+^ T-cells (identified as hCD45^+^CD3^+^CD4^+^ in uninfected mice and hCD45^+^CD3^+^CD8^-^ in HIV-positive mice as HIV downregulates CD4). To analyze T-cells infected with HIV, cells were permeabilized, and intracellular staining with anti-p24 antibody was performed. HIV-infected T-cells were subsequently gated as hCD45^+^CD3^+^CD8^-^p24^+^. Shown is the flow cytometry analysis of spleen cells from a representative HIV-infected CCST mouse.

**Supplemental Figure 5.**
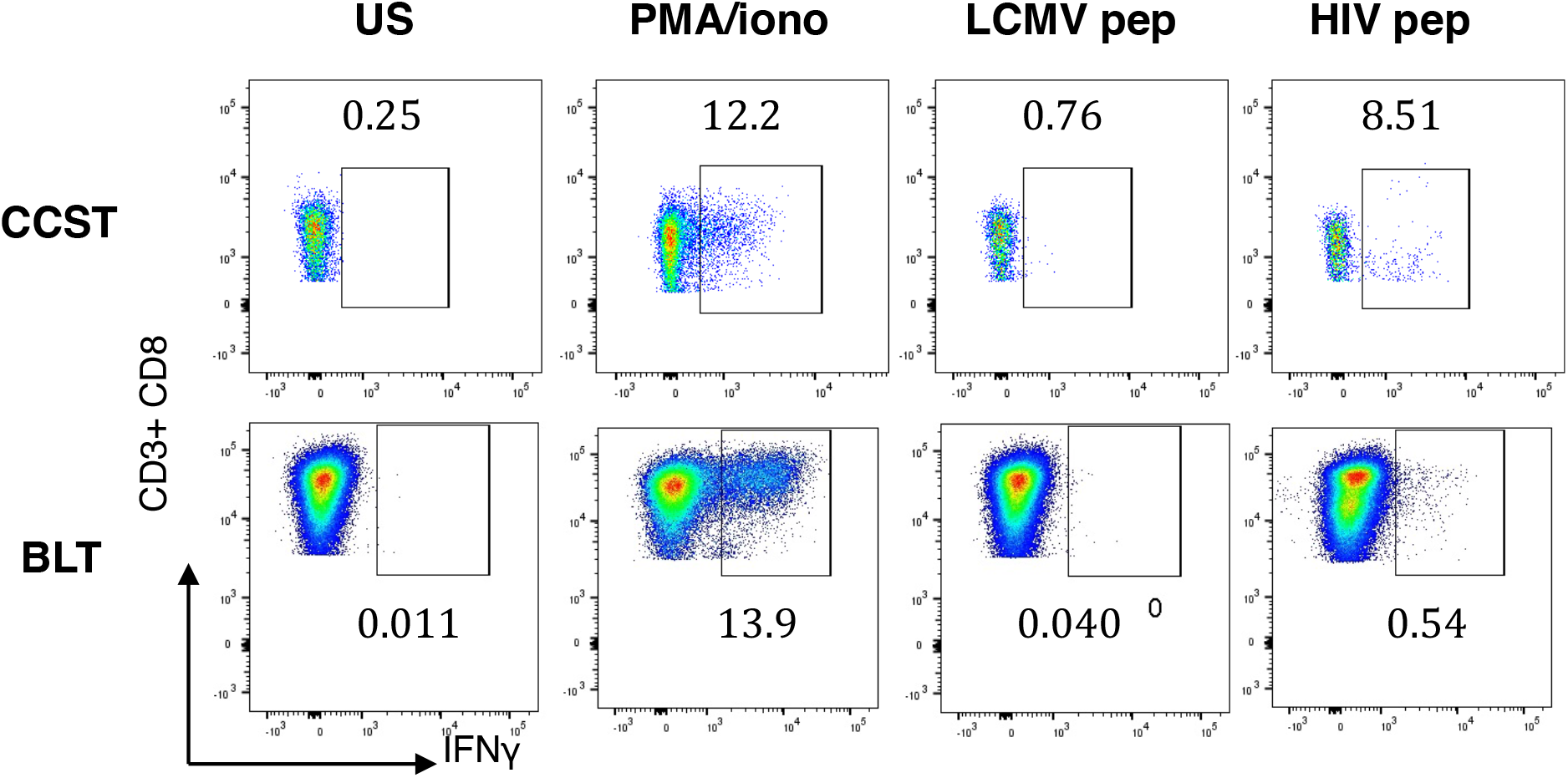
Interferon-γ production by CD8^+^ T cells from CCST and BLT mice upon antigenic stimulation. Splenocytes from infected CCST and BLT mice were kept untreated or stimulated with PMA/ionomycin, pooled Clade B HIV peptides (env, gag, pol, nef) (2 μg/mL) or LCMV peptides (GP61-80, GP276-286) (100 ng/mL). LCMV stimulation served as a specificity control for the assay and frequencies of IFNγ-expressing human T-cells were measured intracellularly by flow cytometry. Shown is an example of cytokine staining in CD8^+^ T cells.

**Supplemental Table 1.** HLA match and immune reconstitution.

|  | Success<br>n (%) | Failure<br>n (%) | Chi-Square |
| --- | --- | --- | --- |
| No HLA match | 3 (100%) | 0 (0%) | 0.9232 |
| 1 match | 56 (95%) | 3 (5%) |  |
| 2 or more matches | 39 (95%) | 2 (5%) |  |

## Notes

### Competing Interest Statement

The authors have declared no competing interest.

